# Establishing the role of SINE proteins in regulating stomatal dynamics in Arabidopsis thaliana

**DOI:** 10.1101/759712

**Authors:** Alecia Biel, Morgan Moser, Iris Meier

## Abstract

Stomatal movement, which regulates gas exchange in plants, is controlled by a variety of environmental factors, including biotic and abiotic stresses. The stress hormone ABA initiates a signaling cascade, which leads to increased H_2_O_2_ and Ca^2+^ levels and F-actin reorganization, but the mechanism of, and connection between, these events is unclear. SINE1, an outer nuclear envelope component of a plant Linker of Nucleoskeleton and Cytoskeleton (LINC) complex, associates with F-actin and is, along with its paralog SINE2, expressed in guard cells. Here, we have determined that Arabidopsis SINE1 and SINE2 play an important role in stomatal regulation. We show that SINE1 and SINE2 are required for stomatal opening and closing. Loss of SINE1 or SINE2 results in ABA hyposensitivity and impaired stomatal dynamics but does not affect stomatal closure induced by the bacterial elicitor flg22. The ABA-induced stomatal closure phenotype is, in part, attributed to impairments in Ca^2+^ and F-actin regulation. Together, the data suggest that SINE1 and SINE2 act downstream of ABA but upstream of Ca^2+^ and F-actin. While there is a large degree of functional overlap between the two proteins, there are also critical differences. Our study makes an unanticipated connection between stomatal regulation and a novel class of nuclear envelope proteins, and adds two new players to the increasingly complex system of guard cell regulation.

## Introduction

Eukaryotic nuclei are double membrane-bound organelles with distinct but continuous inner nuclear membranes (INM) and outer nuclear membranes (ONM). The site where the INM and ONM meet forms the nuclear pore, where nucleocytoplasmic transport occurs (Jevtić et al., 2014). The linker of the nucleoskeleton and cytoskeleton (LINC) complexes are protein complexes spanning the inner and outer nuclear envelope. They contribute to nuclear morphology, nuclear movement and positioning, chromatin organization and gene expression, and have been connected to human diseases (Chang et al., 2015; Chang et al., 2015; Lv et al., 2015). LINC complexes are comprised of <u>K</u>larsicht/<u>A</u>NC-1/<u>S</u>yne <u>H</u>omology (KASH) ONM proteins and <u>S</u>ad1/<u>UN</u>C-84 (SUN) INM proteins that interact in the lumen of the nuclear envelope (NE), thus forming a bridge between the nucleoplasm and the cytoplasm.

Opisthokonts (animals and fungi) and plants have homologous SUN proteins with C-terminal SUN domains located in the NE lumen. However, no proteins with sequence similarity to animal KASH proteins have been discovered in plants and thus much less is known regarding the role of plant LINC complexes (Graumann et al., 2010; Oda and Fukuda, 2011). Within the past few years, studies identifying structurally similar plant KASH protein analogs have caused increased interest in this area (Graumann et al., 2014; Zhou et al., 2014; Zhou et al., 2015). Arabidopsis ONM-localized WPP domain-interacting proteins (WIPs) were the first identified plant analogs of animal KASH proteins, binding the SUN domain of Arabidopsis SUN1 and SUN2 in the NE lumen (Zhou et al., 2012). WIP1, WIP2, and WIP3 form a complex with WPP-interacting tail-anchored proteins (WIT1 and WIT2). Together, they are involved in anchoring the Ran GTPase activating protein RanGAP to the NE (Zhou et al., 2012), in nuclear movement in leaf mesophyll and epidermal cells and root hairs (Zhou et al., 2012; Tamura et al., 2013; Zhou and Meier, 2013; Tamura et al., 2015), and in nuclear movement in pollen tubes (Zhou et al., 2015).

Based on similarity to the SUN-interacting C-terminal tail domain of WIP1-3, additional plant-unique KASH proteins were identified and named SUN domain-interacting NE proteins (SINE1-SINE4 in Arabidopsis) (Zhou et al., 2014). Arabidopsis SINE1 and SINE2 are paralogues and are conserved among land plants. In leaves, SINE1 is exclusively expressed in guard cells and the guard cell developmental lineage, whereas SINE2 is expressed in trichomes, epidermal and mesophyll cells, and only weakly in mature guard cells (Zhou et al., 2014). Both SINE1 and SINE2 are also expressed in seedling roots and share an N-terminal domain with homology to armadillo (ARM). Proteins encoding ARM repeats have been reported to bind actin and act as a protein-protein interaction domain in a multitude of proteins across both plant and animal kingdoms (Coates, 2003). SINE1 was verified to associate with filamentous actin (F-actin) via its ARM domain through colocalization studies in *N. benthamiana* leaves and Arabidopsis roots but SINE2 does not share this property. Furthermore, depolymerization of F-actin by LatB disrupts GFP-SINE1 localization in guard cells and increases GFP-SINE1 mobility during FRAP analysis, suggesting a SINE1-F-actin interaction in guard cells. Mutant analysis showed that SINE1 is required for the symmetric, paired localization of nuclei in guard cells, while SINE2 contributes to plant immunity against the oomycete pathogen *Hyaloperonospora arabidopsis* (Zhou et al., 2014).

Stomatal dynamics rely on highly coordinated and controlled influx and efflux of water and ions which increase turgor pressure to facilitate opening and decrease turgor for stomatal closing. This process is mediated through complex signal transduction pathways, being controlled by plant and environmental parameters such as changes in light conditions and abiotic and biotic stresses (Schroeder et al., 2001). Light changes result in a conditioned stomatal response in which stomata open and close in a daily cyclic fashion. Abiotic stresses, such as drought, and biotic stresses, such as pathogen exposure, can both override this daily cycle to induce a specific stomatal response.

The plant hormone abscisic acid (ABA) senses and responds to abiotic stresses, with ABA metabolic enzymes regulated by changes in drought, salinity, temperature, and light (Zhang et al., 2008; Xi et al., 2010; Verma et al., 2016). ABA initiates long-term responses, such as growth regulation, through alterations in gene expression (Kang et al., 2002; Fujita et al., 2005) and induces stomatal closure as a short-term response to stress, involving the activation of guard cell anion channels and cytoskeleton reorganization (Eun and Lee, 1997; Zhao et al., 2011; Jiang et al., 2012; Li et al., 2014). F-actin is radially arrayed in open guard cells of several diverse plant species and undergoes reorganization into a linear or diffuse bundled array upon stomatal closure (Kim et al., 1995; Xiao et al., 2004; Li et al., 2014; Zhao et al., 2016). Although many disparate players have been shown to be important for regulating stomatal dynamics, it is still unclear how these events are interconnected and where actin reorganization fits in.

Here, we have investigated if Arabidopsis SINE1 and SINE2 play a physiological role in guard cell biology. Our findings show that both SINE1 and SINE2 are required for stomatal opening and closing. Loss of SINE1 or SINE2 results in ABA hyposensitivity and impaired stomatal dynamics but does not affect pathogen-induced stomatal closure from the bacterial peptide flg22. The ABA-induced stomatal closure phenotype is, in part, attributed to impairments in calcium and actin regulation.

## Results

### SINE1 and SINE2 are involved in light regulation of stomatal opening and closing

To assess whether SINE1 and SINE2 have a function in guard cell dynamics, we first monitored stomatal aperture changes in *sine1-1, sine2-1,* and *sine1-1 sine2-1* double mutants when exposed to light-dark cycles using *in vivo* stomatal imprints from attached leaves. At the start of the assay, two hours before lights were turned on, average stomatal apertures were between 2.8 µm and 3.3 µm (Fig. 1A). By mid-day, four hours after the lights were turned on, WT stomata were fully opened, while *sine1-1* and *sine2-1* mutant stomata had opened only marginally. Expression of proSINE1:GFP-SINE1 in *sine1-1* (SINE1:*sine1-1*) or proSINE2:GFP-SINE2 in *sine2-1* (SINE2:*sine2-1*) partially restored stomatal responsiveness to changes in light conditions, whereas the *sine1-1 sine2-1* double mutant plants displayed intermediate changes in stomatal dynamics. On average, neither *sine1-1* nor *sine2-1* mutant stomata were fully open or fully closed for the duration of the assay.

**Figure 1:**
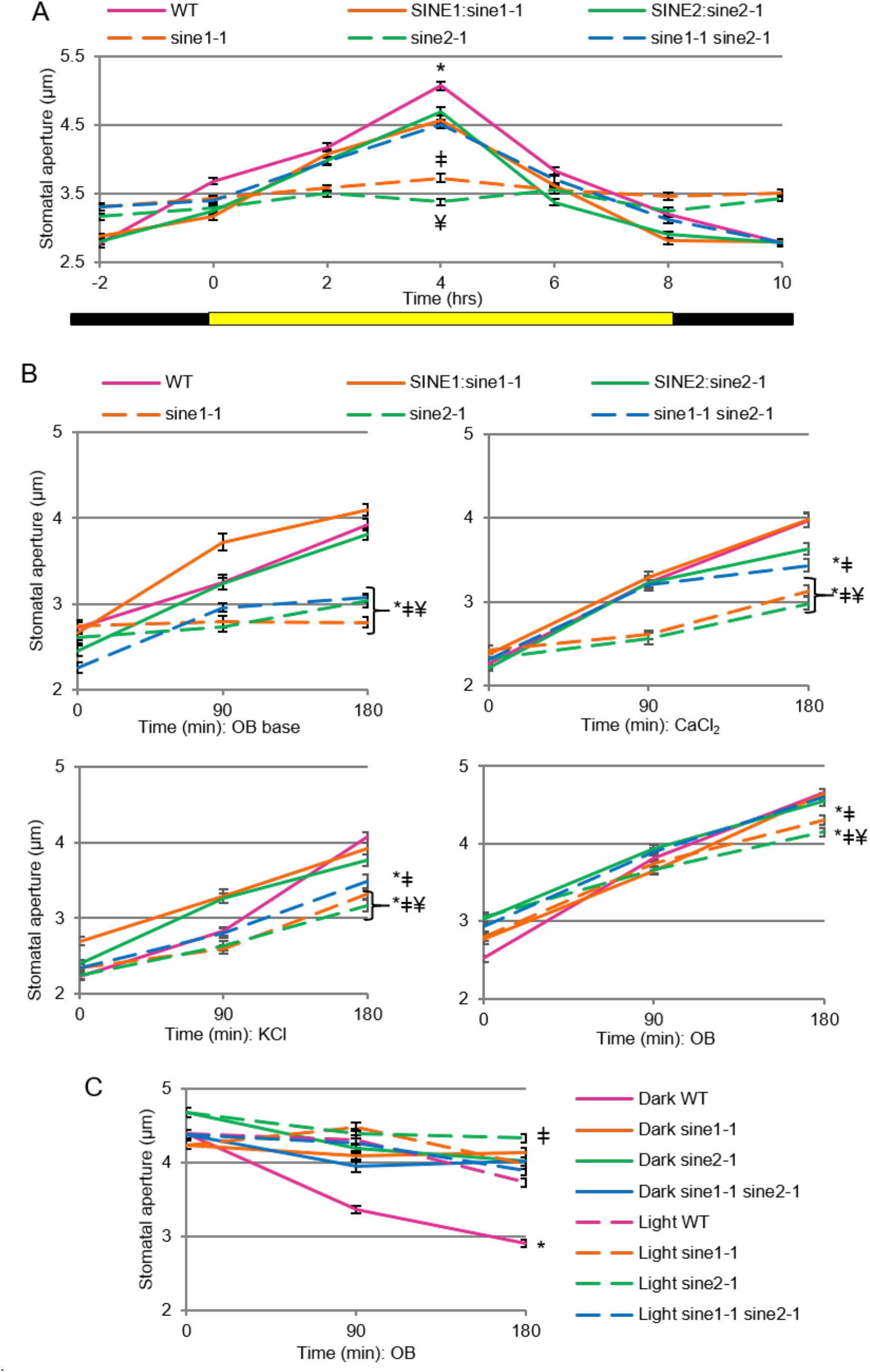
Determining the role of SINE1 and SINE2 in the light regulation of stomatal dynamics. (A) Stomatal imprints from intact whole Arabidopsis leaves were taken and stomatal apertures were measured 2 h prior to the onset of lights (yellow bar) and every 2 h thereafter until 2 h after lights off (black bar). Symbols denote statistical significance, with P<0.001. *: WT vs. all other lines; ǂ: *sine1-1* vs. WT, SINE1:*sine1-1*, and *sine1-1 sine2-1*; ¥: *sine 2-1* vs. WT, SINE2:*sine2-1*, and *sine1-1 sine2-1*; (B) Whole leaves were placed in specified buffer for 3 h under constant light at the end of a night cycle, epidermal peels were mounted every 90 min, and stomatal apertures were measured. Top-left: opening buffer (OB) base (see methods); Top-right: OB base plus 10 µM CaCl_2_; Bottom-left: OB base plus 20 mM KCl; Bottom-right: OB base plus 10 µM CaCl_2_ and 20 mM KCl. Symbols denote statistical significance, with P<0.001. *: specified lines vs WT; ǂ: specified lines vs. SINE1:*sine1-1*; ¥: specified lines vs. SINE2:*sine2-1*; (C) Whole leaves were placed in OB under constant light for 3 h and either kept under constant light or placed in dark for an additional 3 h. Epidermal peels were taken and stomatal apertures were measured every 90 min after the initial 3 h stomatal opening phase. Symbols denote statistical significance as determined by Student’s t-test, with P<0.001. *: dark WT vs light WT; ǂ: dark *sine2-1* vs light *sine2-1*. All data are mean values ± SE from three independent experiments.

To further assess stomatal opening, detached leaves from WT and *sine* mutants (*sine1-1*, *sine2-1*, and *sine1-1 sine2-1*) were incubated in buffers containing Ca^2+^, K^+^, Ca^2+^ and K^+^, or neither ion (Fig. 1B). In the absence of external K^+^ and Ca^2+^ (opening buffer (OB) base), light-induced stomatal opening was impaired in *sine1-1*, *sine2-1*, and *sine1-1 sine2-1* (Fig. 1B, top left panel). With exposure to external Ca^2+^ (20 µM CaCl_2_), *sine1-1*, *sine2-1*, and *sine1-1 sine2-1* still displayed significantly impaired stomatal opening (Fig. 1B, top right panel). Likewise, with exposure to external K^+^ (50 mM KCl), statistically significant impairment during opening was seen in both single and double mutants compared to WT (Fig. 1B, bottom left panel). When leaves were exposed to both Ca^2+^ and K^+^ (OB), stomatal opening in *sine1-1* and *sine2-1* was still somewhat reduced (Fig. 1B, bottom right panel), however, opening was greatly increased compared to the conditions lacking one or both ions. Similar positive effects of low concentrations of Ca^2+^ on stomatal opening have been previously described (Hao et al., 2012; Wang et al., 2014). Under these conditions (OB incubation), the double mutant behaved similar to WT. In all assays, SINE1:*sine1-1* and SINE2:*sine2-1* displayed similar opening to WT, indicating full rescue of the mutations in these lines. Finally, stomatal closure was assessed after leaves were either transitioned from 3 hours of light to 3 hours of dark or kept under constant light. *sine1-1*, *sine2-1*, and *sine1-1 sine2-1* were significantly impaired in closing response, suggesting that SINE1 and SINE2 are also required for dark-induced stomatal closure (Fig. 1C). Together, these data indicate that SINE1 and SINE2 are involved in stomatal opening in response to white light and in closing in response to dark, and that exogenous Ca^2+^ and K^+^ can at least partially rescue the opening defect.

### Impaired ABA-induced stomatal closure in *sine1-1* and *sine2-1*

Abscisic acid (ABA) has been widely used to induce stomatal closure and monitor stomatal response to simulated abiotic stress (Umezawa et al., 2010) and was used here to test ABA stomatal response in *sine* mutants. Prior to this assay, we tested stomatal opening for all lines used here to ensure equal starting conditions for the closing assays (Supplemental Fig. 1).

A difference in stomatal aperture was noticed as early as one hour after addition of 20 µM ABA in all *sine* mutants when compared to WT (Fig. 2A). WT stomata continued to close over the following two hours, while *sine1-1* (Fig. 2A, left panel), *sine2-1* (Fig. 2A, right panel), and *sine1-1 sine2-1* (Fig. 2A left and right panel) did not exhibit further stomatal closure. All data in Fig. 2A were collected at the same time but are split here into two panels for presentation purposes. WT and *sine1-1 sine2-1* traces are therefore shown twice. In this assay, *sine1-1 sine2-1* stomatal closure resembled that of *sine1-1* and *sine2-1* single mutants. This suggests that there is no additive effect of the *sine1-1* and *sine2-1* mutants in this response, indicating that SINE1 and SINE2 are working in the same pathway. Figure 2B shows representative images of WT and *sine1-1* stomatal apertures before and after three hours of exposure to ABA.

**Figure 2:**
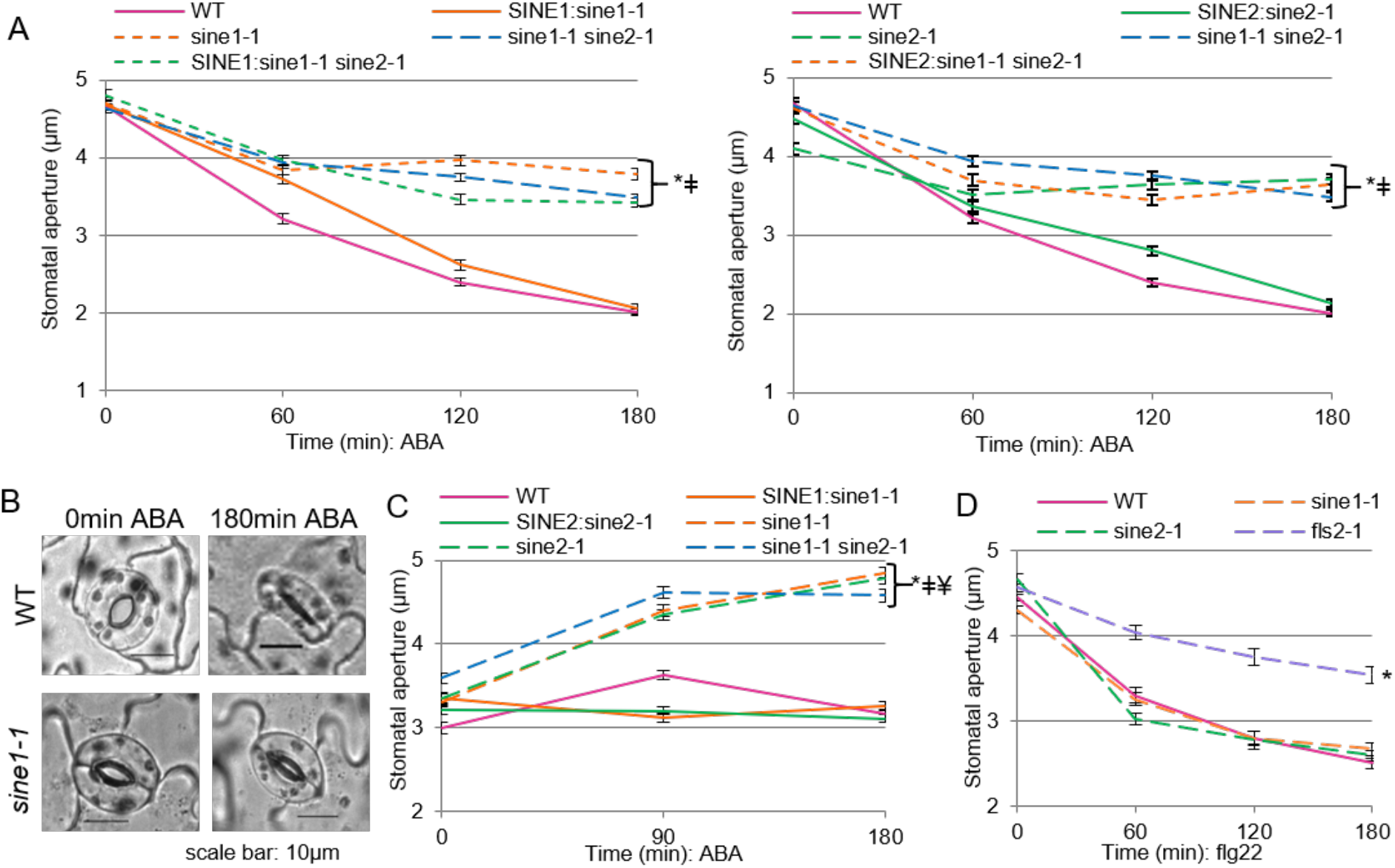
Abiotic vs. biotic stress induced stomatal changes in sine mutants. Stomatal opening and closing assays were used here as described in methods. (A) leaves incubated in 20 µM ABA during closure; Data obtained from one experiment and split into two panels for clarity (B) Representative images of stomata for WT and *sine1-1* lines before and after ABA exposure. (C) leaves were incubated in OB plus 20 µM ABA during stomatal opening. (D) leaves incubated in 5 µM flg22 during closure. All data are mean values ± SE from three independent experiments. Symbols denote statistical significance as determined by Student’s t-test, with P<0.001. *: specified lines vs. WT; ǂ: specified lines vs. SINE1:*sine1-1*; ¥: specified lines vs. SINE2:*sine2-1*.

Exogenous ABA induced stomatal closure in SINE1:*sine1-1* and SINE2:*sine2-1* at a similar rate as in WT (Fig. 2A). To further verify the importance of both SINE1 and SINE2 in ABA-induced stomatal closure, the following ‘partially’ complemented lines were used: SINE1pro:GFP-SINE1 in *sine1-1 sine2-1* (SINE1:*sine1-1 sine2-1*), and SINE2pro:GFP-SINE2 in *sine1-1 sine2-1* (SINE2:*sine1-1 sine2-1*) (Zhou et al., 2014). Upon ABA exposure, SINE1:*sine1-1 sine2-1* and SINE2:*sine1-1 sine2-1* showed significantly impaired stomatal closure compared to WT. Hence, neither SINE1 nor SINE2 alone is sufficient to rescue the *sine1-1 sine2-1* phenotype, confirming the single and double mutant analysis.

In order to verify that this phenotype holds true for multiple alleles, the additional T-DNA insertion alleles *sine1-3* and *sine2-2* were used to assay single and double mutant lines along with the SINE1:*sine1-3* and SINE2:*sine2-2* complemented lines and the ‘partially’ complemented lines SINE1:*sine1-3 sine2-1* and SINE2:*sine1-3 sine2-2* (Zhou et al., 2014). The same ABA-induced stomatal closure assay was performed as seen in Figure 2A and similar results were obtained, confirming independence of the phenotypes from insertion position and genetic background (Supplemental Fig. 2). Therefore, only the *sine1-1* and *sine2-1* mutants were used for the subsequent assays.

In addition to inducing stomatal closure, ABA also inhibits stomatal opening (Yin et al., 2013). To test the role of SINE1 and SINE2 in this process, stomatal opening assays utilizing OB were performed in the presence of 20 µM ABA (Fig. 2C). Under these conditions, WT, SINE1:*sine1-1*, and SINE2:*sine2-1* are unable to open stomata in the presence of ABA. However, stomata of *sine1-1*, *sine2-1*, and *sine1-1 sine2-1* are unimpeded in their ability to open, indicating that loss of SINE1 or SINE2 results in impaired stomatal response to ABA for both opening and closing.

Stomatal closure can also be induced by biotic stresses, such as pathogen exposure (Zhang et al., 2008; Guzel Deger et al., 2015). The bacterial elicitor flg22 is a well-studied and accepted tool to simulate pathogen-induced stomatal closure, and was therefore tested here. As a control, we used the LRR receptor-like kinase mutant *fls2-1*, which is unable to recognize and bind flg22 (Dunning et al., 2007). After exposure to 5 µM flg22, *fls2-1* had significantly inhibited stomatal closure compared to WT, while flg22-induced stomatal closure in *sine1-1* and *sine2-1* was similar to that of WT (Fig. 2D). Together, these data indicate that SINE1 and SINE2 are working within the ABA pathway to regulate ABA-induced stomatal closure and ABA-inhibition of stomatal opening and that these roles are distinct from the flg22-induced stomatal closure pathway.

### Drought susceptibility is increased in *sine1-1* and *sine2-1* plants after stomatal opening

We next wanted to determine if the impaired stomatal dynamics observed for *sine1-1* and *sine2-1* are detrimental to plant vitality. If stomata are unable to close in response to stress, it is expected that an increase in transpiration would occur (Kang et al., 2002; Mustilli et al., 2002). As an initial investigation into drought susceptibility, we measured weight loss of freshly detached leaves at midday. Detached leaves were kept in a petri dish and weighed collectively for each genotype. Fresh weight loss was similar in *sine2-1*, *sine1-1 sine2-1*, SINE1:*sine1-1*, and SINE2:*sine2-1* compared to WT (Fig. 3A). Although *sine1-1* did show a statistically significant increase in fresh weight loss compared to WT (P<0.05), there was no difference observed between *sine1-1* and SINE1:*sine1-1*. Thus, despite the stomatal dynamics phenotypes reported above, there appears to be no conclusive difference in fresh weight loss between WT, *sine1-1* and *sine2-1* freshly detached leaves. We reasoned that this might be due to the fact that *sine1-1* and *sine2-1* were impaired both in opening and closing (Fig. 1A), thus likely leading to only semi-open stomata in the detached leaves of the *sine1-1* and *sine2-1* mutants.

**Figure 3:**
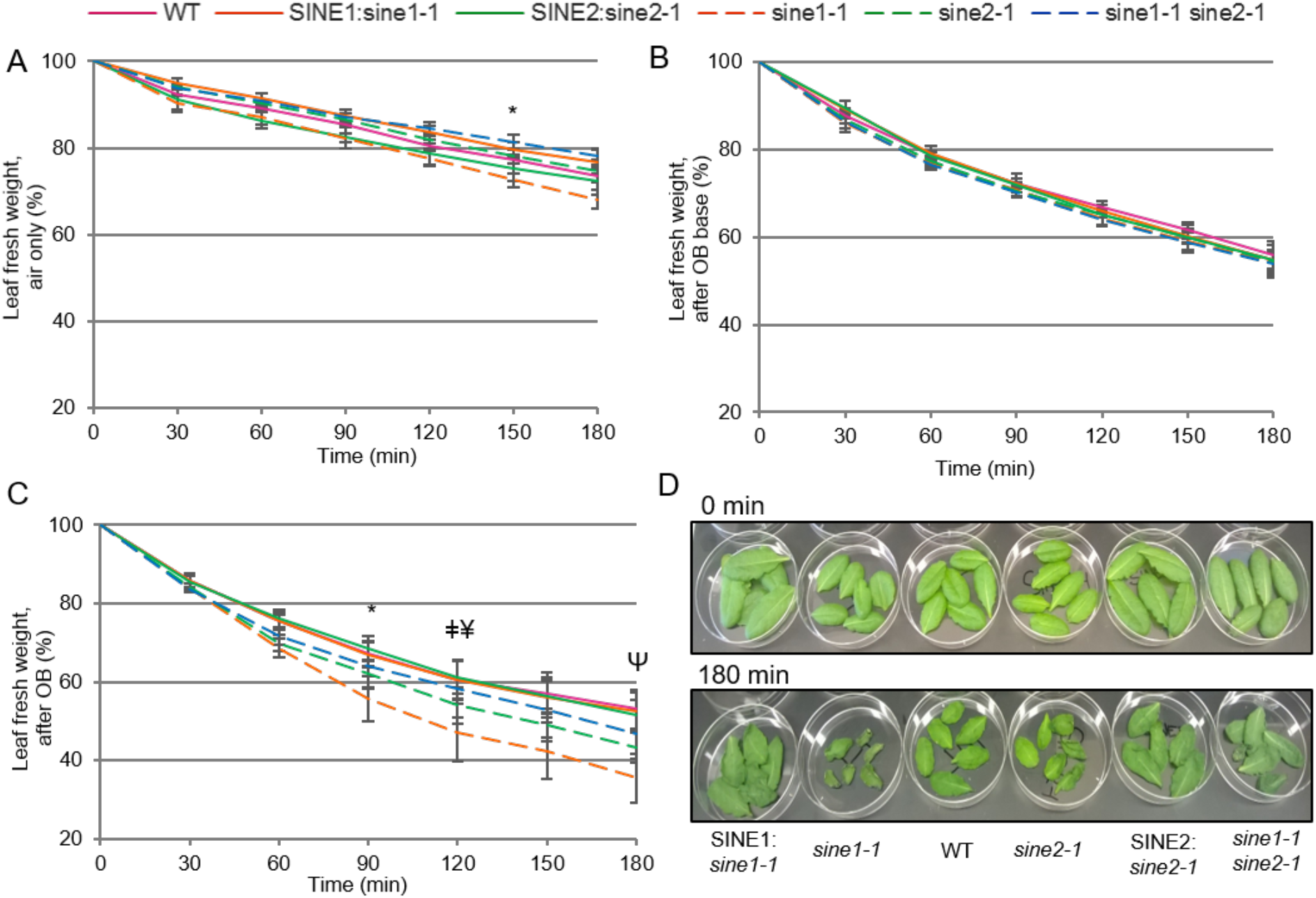
Transpiration rates after induced stomatal opening. Rosette leaves were taken from 6-8 week old short day plants at similar developmental stages for each of the lines depicted and kept abaxial side up. Fresh leaves were placed in specified buffer for 3 hours under constant light, transferred to a petri dish as specified below, and weighed every 30 minutes thereafter. (A) in air; (B) in opening buffer (OB) base; (C) in OB with representative images shown in (D) where the top images are leaves at 0 minutes before OB incubation and bottom images are leaves after 180 min OB incubation. Mean values ± SE from at least three independent experiments are shown in A-C. (B) shows no statistically significant differences. Symbols in (A) and (C) denote the beginning of statistically significant differences as determined by Student’s t-test, with P<0.05. *: *sine1-1* vs. WT; ǂ: *sine2-1* vs. WT; ¥: *sine2-1* vs. SINE2:*sine2-1*; d: *sine1-1* vs. SINE1:*sine1-1*.

To test this hypothesis, we repeated the water loss assay, but incubated leaves for 3 hours in OB before transferring them to air and monitoring fresh weight loss. As a control, detached leaves of *sine1-1*, *sine2-1*, and *sine1-1 sine2-1* were exposed to OB base (without calcium or potassium, Fig. 3B) and similar results were obtained as seen in Figure 3*: no differences were observed in leaf fresh weight loss between the tested lines. However, after pre-exposure to OB for three hours, *sine1-1* and *sine2-1* leaves lost weight at a significantly faster rate than WT while SINE1:*sine1-1* and SINE2:*sine2-1* lost weight at similar rates as WT and *sine1-1 sine2-1* showed an intermediate phenotype with no statistically significant difference to WT (Fig. 3C, P<0.05). Leaf morphology of *sine1-1* and *sine2-1* agreed with these observations in that pre-exposure to OB led to more rapid wilting, as seen by increased leaf curling and shrinking compared to WT (Fig. 3D).

The water loss assay was also repeated with a slightly different experimental setting: First, individual leaves were placed abaxially side up throughout the assay and weighed separately to avoid a potential influence of overlapping leaves (Supplemental Fig. S3A-B), and second, an additional T-DNA allele combination for the double mutant (*sine1-3 sine2-2*) was added to exclude a potential influence of the genetic background. In addition, two lines expressing SINE1 and SINE2 under control of the 35S promoter in a WT background were added: 35S:GFP-SINE1 in WT (SINE1:WT) and 35S:GFP-SINE1 in WT (SINE2:WT). The data from this assay largely recapitulated those shown in Figure 3, suggesting that the different incubation conditions did not influence the assay. In addition, both double mutants lost water at an intermediate rate, compared to WT and the single mutants (Supplemental Fig. S3C), again consistent with the leaf morphology at the end of the assay (Supplemental Fig. S3B). In combination with the stomatal opening dynamics phenotypes seen in Figure 1, these data indicate that under the conditions assayed in Figure 3A, no increased susceptibility to drought was seen, likely because the opening and closing defects cancel each other out. However, after forced stomatal opening (compare Fig. 1B last panel), increased drought susceptibility was revealed in *sine1-1*, *sine2-1*, and partially in *sine1-1 sine2-1*, consistent with the altered stomatal dynamics.

### Mapping the position of SINE1 and SINE2 in the stomatal ABA signaling pathway

Upon ABA perception, a signaling cascade results in the induction of both H_2_O_2_ and Ca^2+^ (Pei et al., 2000; Umezawa et al., 2010; Zhao et al., 2011). To narrow down the position of SINE1 and SINE2 in ABA-induced stomatal closure, we therefore investigated hydrogen peroxide-induced and calcium-induced stomatal closure. Stomatal closure was measured as described above, in response to either 0.5 mM H_2_O_2_ or 2 mM CaCl_2_ (Zhao et al., 2011). Upon exposure to H_2_O_2_, stomatal closure was impaired in *sine1-1*, *sine2-1* and *sine1-1 sine2-1* compared to WT (Fig. 4A, 4B). SINE1:*sine1-1* and SINE2:*sine2-1* displayed H_2_O_2_-induced stomatal closure similar to WT (Fig. 4A, 4B). Significantly reduced stomatal closure was seen in SINE1:*sine1-1 sine2-1* and SINE2:*sine1-1 sine2-1*, again confirming the single and double mutant results (Fig. 4A, 4B).

**Figure 4:**
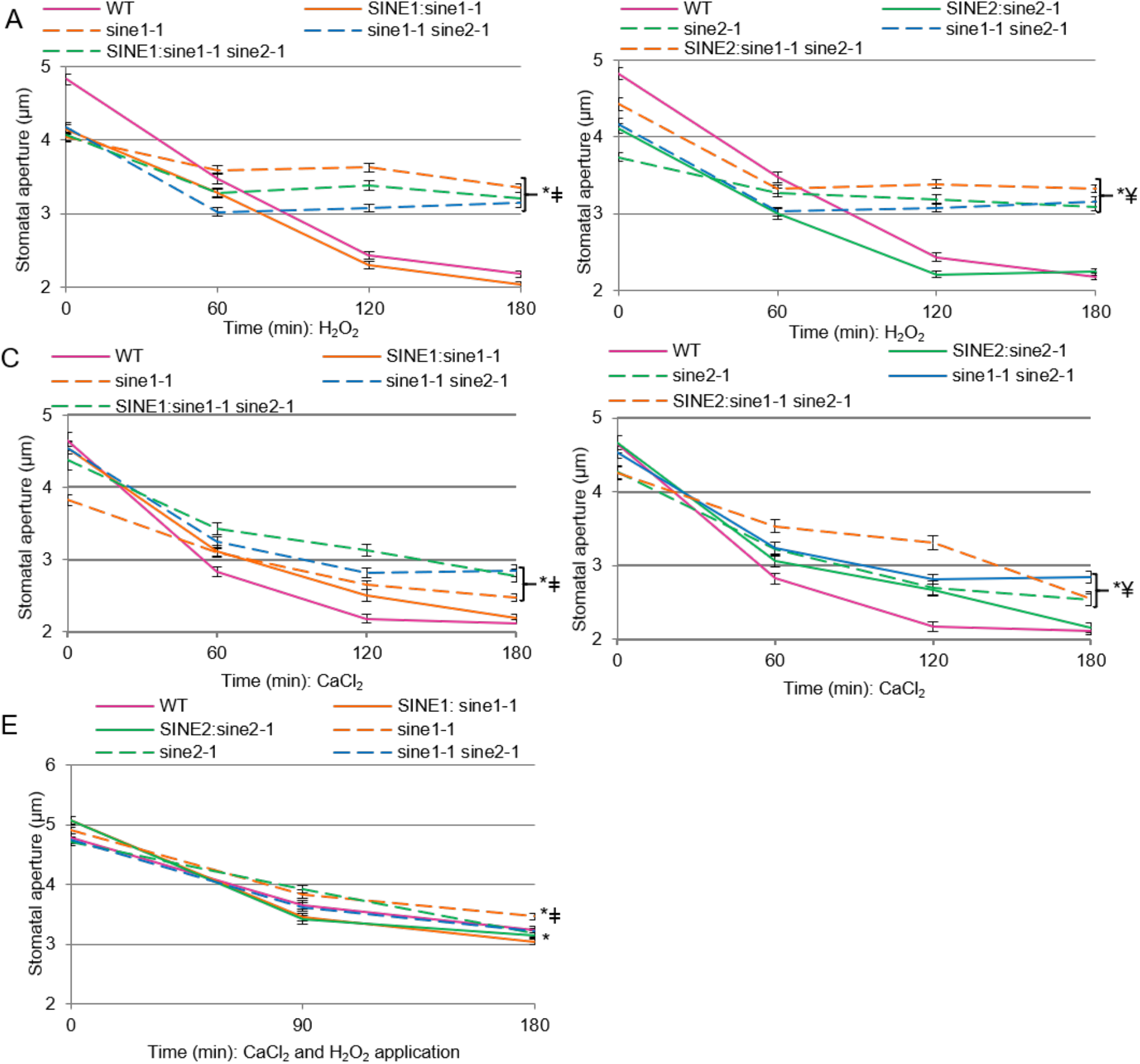
Stomatal closure in response to H_2_O_2_ and CaCl_2_ for SINE1 and SINE2 mutants. Stomatal opening and closing assays were used here as described in methods. (A-B) leaves incubated in 0.5 mM H_2_O_2_ during closure; Data obtained from one experiment and split into two panels for clarity (C-D) leaves incubated in 2 mM CaCl_2_ during closure; Data obtained from one experiment and split into two panels for clarity. (E) leaves incubated in 0.5 mM H_2_O_2_ plus 2 mM CaCl_2_ during closure; All data are mean values ± SE from three independent experiments. Symbols denote statistical significance as determined by Student’s t-test, with P<0.001. *: specified lines vs. WT; ǂ: specified lines vs. GFP-SINE1:*sine1-1*; ¥: specified lines vs. GFP-SINE2:*sine2-1*.

When exposed to a Ca^2+^ donor, CaCl_2_, *sine1-1*, *sine2-1,* and *sine1-1 sine2-1* were somewhat impaired in stomatal closure, which was also observed in the double mutant (Fig. 4C, 4D). (As above, data shown were obtained at the same time and split for clarification.) However, 2mM CaCl_2_ more effectively triggered stomatal closure in *sine1-1, sine2-1,* and *sine1-1 sine2-1* than the previous treatments of ABA or H_2_O_2_ (Supplemental Table S1). These results indicate that external calcium is able to partially rescue the stomatal closure phenotype. Meanwhile, SINE1:*sine1-1* and SINE2:*sine2-1* lines showed stomatal closure similar to WT in response to exogenous application of calcium and SINE1:*sine1-1 sine2-1* and SINE2:*sine1-1 sine2-1* had a similar degree of stomatal closure as single and double *sine* mutants (Fig. 4C, 4D). Together, these data show that the impaired stomatal closure response of SINE1 and SINE2 mutants can be partially rescued by Ca^2+^, but not by H_2_O_2_.

Within the ABA pathway, there is feedback between the Ca^2+^ and H_2_O_2_ branches (Pei et al., 2000; Desikan et al., 2004; Zou et al., 2015). Thus, we also tested the stomatal response of *sine1-1* and *sine2-1* to a combination of both inducers. With exposure to both Ca^2+^ and H_2_O_2_, stomatal closure was similar between *sine2-1*, *sine1-1 sine2-1*, WT, SINE1:*sine1-1*, and SINE2:*sine2-1* (Fig. 4E). Although the stomata of *sine1-1* mutants were statistically more open compared to WT (Fig. 4E, P<0.001), this was by a very small difference (3.46µm vs. 3.15µm, respectively). Finally, ABA-, H_2_O_2_-, Ca^2+^-, and darkness-induced stomatal closure was compared as percent closure to rule out bias introduced by possibly different apertures at the beginning of each assay (Supplemental Table S2; see Materials and Methods). This did not lead to any change in the data interpretation described above.

### Stomatal overexpression of SINE2 leads to compromised stomatal dynamics

Thus far, loss of either *sine1-1* or *sine2-1* has been shown to compromise stomatal dynamics in a similar manner. As previously mentioned, SINE1 and SINE2 show different levels of endogenous protein expression as well as different expression patterns. Thus, we assessed the impact of ubiquitous expression of these proteins on stomatal opening and closing. 35S:GFP-SINE1 in WT (SINE1:WT) and 35S:GFP-SINE2 in WT (SINE2:WT), respectively, were compared to SINE1pro:GFP-SINE1 (SINE1:*sine1-1*) and SINE2pro:GFP-SINE2 (SINE2:*sine2-1*). Confocal microscopy showed that SINE1:WT and SINE1:*sine1-1* have similar expression levels in guard cells (Fig. 5A, top panels). However, as expected, SINE2:WT showed significantly higher GFP expression in guard cells than SINE2:*sine2-1*. Indeed, under the assay conditions, no GFP signal above background was detected in SINE2:*sine2-1* expressing guard cells (Fig. 5A, bottom panels). (A faint nuclear envelope signal was detectable in SINE2:*sine2-1* with higher gain and laser settings, see Materials and Methods.) This observation was further verified by quantifying the nucleus-associated fluorescent signal (Fig. 5B). In contrast, immunoblots of protein extracts from whole seedlings and rosette leaves of SINE1:WT, SINE2:WT, SINE1:*sine1-1* and SINE2:*sine2-1* showed similar amounts of GFP-fusion protein (Supplemental Fig. S4). This confirms GFP-SINE2 expression in SINE2:*sine2-1* and indicates that SINE2:WT leads to overexpression of GFP-SINE2 in guard cells compared to the native SINE2 promoter.

**Figure 5:**
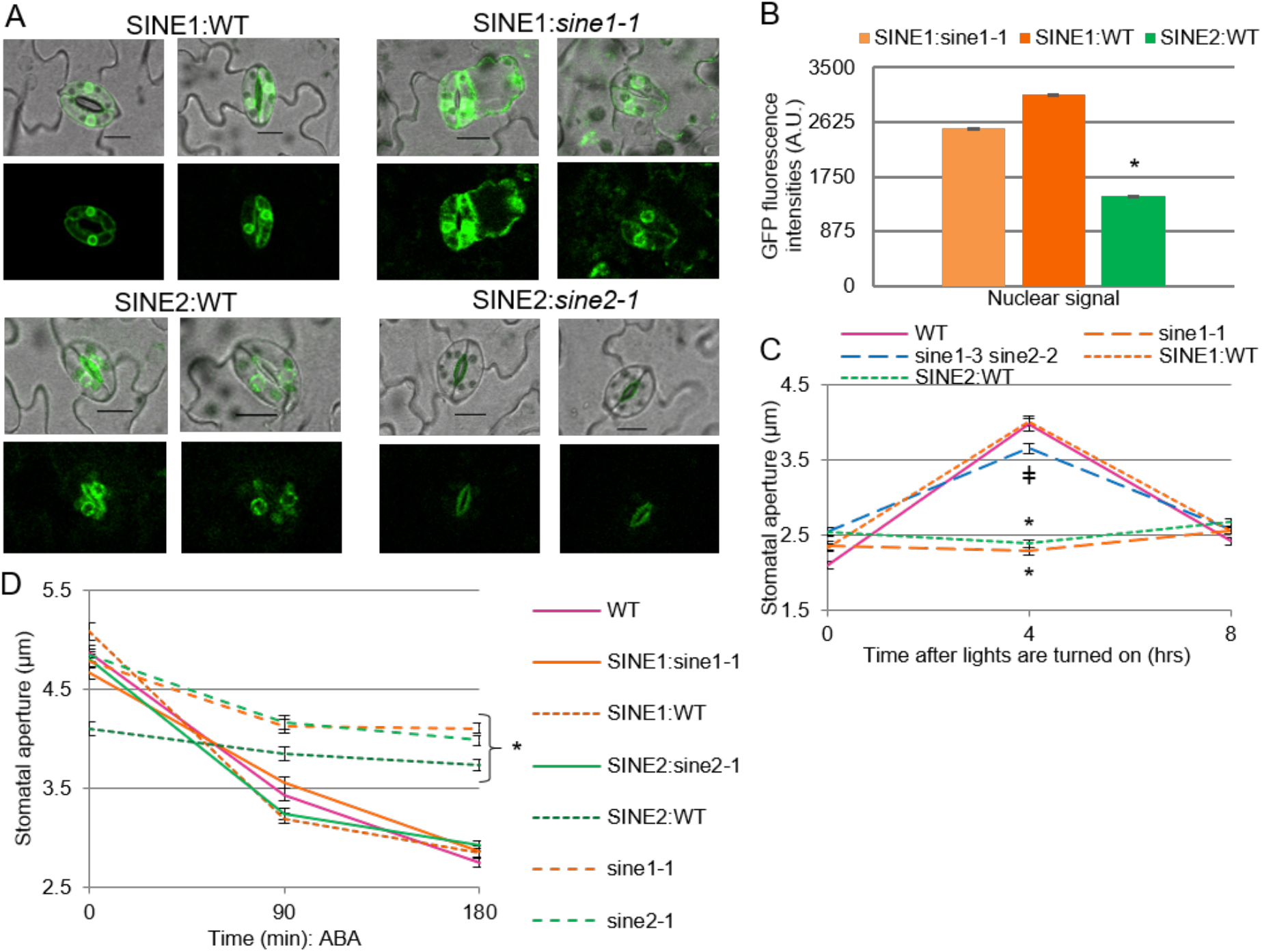
SINE2 but not SINE1 overexpression leads to compromised stomatal dynamics. (A) Confocal microscopy was used to take images of plants expressing GFP-tagged SINE1 or SINE2, with representative images shown. Gain was set to first image of top row and used for the remaining images. Scale bar represents 10 µm. (B) Data taken from two plants. ≥ 50 nuclei were measured for each line. Nuclear fluorescence intensities were measured using ImageJ. SINE2:*sine2-1* had no measurable fluorescence signal at the nucleus and was therefore not quantified. Symbols denote statistical significance as determined by Student’s t-test. *: P<0.005, SINE2:WT vs. SINE1:*sine1-1*. (C) Stomatal apertures taken from stomatal imprint assay at the start and end of light cycles. *: P<0.001, specified lines vs. WT; ǂ: P=0.004, specified lines vs. WT. (D) ABA induced stomatal closure assay as described in methods. *: P<0.001, specified lines vs. WT, SINE1:sine1-1, SINE1:WT, and SINE2:sine2-1. Data shown in (C) and (D) are from three independent experiments. Data are mean values ± SE. Symbols denote statistical significance.

Loss of SINE1 or SINE2 resulted in impairments in stomatal dynamics by both light and dark (Fig. 1A) and ABA (Fig. 2A). We therefore used these two assays to also test stomatal impairments in the SINE1 and SINE2 ubiquitously expressing lines. While SINE1:WT behaved like WT, SINE2:WT recapitulated the *sine1-1* phenotype during a light/dark cycle (Fig. 5C). Similarly, SINE2:WT was largely unresponsive to ABA, while SINE1:WT showed WT-like stomatal closure in response to ABA (Fig. 5D). These data suggest that 35S promoter-driven GFP-SINE1 expression has no significant effect on SINE1/SINE2 function in guard cells. However, additional expression of the normally lowly expressed SINE2 in guard cells appears toxic to SINE1/SINE2 function. This suggests that fine-tuning of cellular abundance of the two proteins is required for their function. Because one model to account for the interference of SINE2 is that accumulation of a SINE1/SINE2 heterodimer could negatively affect a specific role of SINE1 in guard cells, we tested if the two proteins can interact in a split-ubiquitin yeast two-hybrid assay. Indeed, interaction was seen between SINE1 and SINE2 as well as a weaker interaction between SINE2 and SINE2. Because of self-activation issues, the SINE1-SINE1 interaction could not be tested (Supplemental Fig. S5).

Although SINE2:WT recapitulates the *sine1-1* and *sine2-1* phenotype in both a light/dark cycle and in ABA response, this line showed WT-like loss of fresh weight during desiccation (Supplemental Fig. S3). This could be explained if SINE2:WT had a compensatory phenotype, such as altered stomatal density. SINE2 is normally expressed only in mature guard cells whereas SINE1 is expressed in both progenitor guard cells and mature guard cells (Zhou et al., 2014). We therefore tested if 35S promoter-driven SINE2 is also influencing stomatal development (Lucas et al., 2006; Nadeau and Sack, 2002). Indeed, both stomatal index (SI) and stomatal density (SD) are reduced in SINE2:WT, but not in SINE1:WT, WT, or the T-DNA insertion mutants, suggesting that a compensatory phenotype might indeed exist (Supplemental Fig S6).

Together, these data show that ubiquitous expression of SINE2 impairs stomatal response to changes in light conditions as well as during ABA-induced stomatal closure. Additionally, ubiquitous SINE2 expression resulted in altered stomatal development.

### Interactions between *sine1-1* and *sine2-1* mutants and the actin cytoskeleton

F-actin rearrangement has been implicated in stomatal dynamics and undergoes a specific pattern of reorganization (Staiger et al., 2009). When this actin rearrangement is disrupted, there are concomitant perturbations in stomatal dynamics (Kim et al., 1995; Xiao et al., 2004; Jiang et al., 2012; Li et al., 2014; Zhao et al., 2016). We tested here if the characterized *sine* mutants showed interactions with drug-induced F-actin depolymerization or with F-actin stabilization during stomatal closing. Latrunculin B (LatB) results in F-actin depolymerization and facilitates stomatal closure when in the presence of ABA (MacRobbie and Kurup, 2007). In contrast, jasplakinolide (JK) stabilizes and polymerizes F-actin and favors open stomata, inhibiting stomatal closure (MacRobbie and Kurup, 2007; Li et al., 2014). We used LatB and JK in the presence and absence of ABA to assess their influence on *sine1-1* and *sine2-1* stomatal closure.

When WT leaves are incubated in either OB or OB + LatB in the light, stomatal apertures remained open during the three-hour assay (Fig. 6A). Similarly, both OB and OB + LatB exposure resulted in stomata that remained open throughout the assay for *sine1-1* (Fig. 6A, left panel) and *sine2-1* (Fig. 6A, right panel). ABA alone and ABA + LatB were both able to induce stomatal closure in WT, as reported previously (MacRobbie and Kurup, 2007; Fig. 6A). ABA exposure in *sine1-1* and *sine2-1* resulted in minimal closure, as was seen in the previous assays. However, the combination of ABA and LatB resulted in significant closure of stomata in both *sine1-1* (Fig. 6A, left panel P<0.001) and *sine2-1* (Fig. 6A, right panel P<0.001), closely resembling WT. This suggests that LatB treatment overcomes the inhibition of stomatal closure caused by the loss of either SINE1 or SINE2.

**Figure 6:**
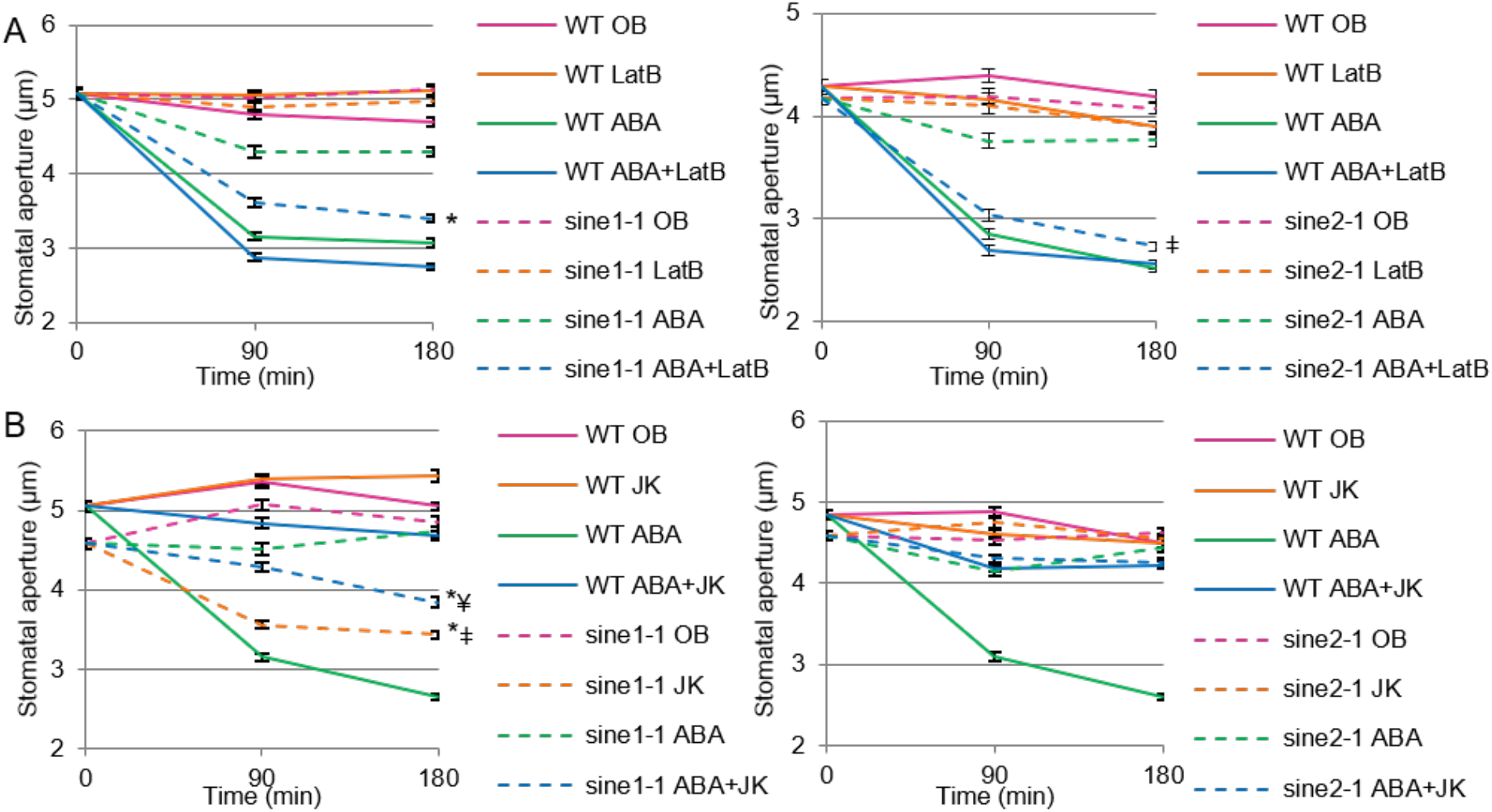
Disrupting actin dynamics in *sine1-1* and *sine2-1* mutant lines alters stomatal closure. Stomatal closure was monitored over a 3 h incubation time in the presence and absence of ABA and actin disrupting drugs. (A) Buffers with and without 20 µM ABA and 10 µM of the F-actin depolymerizing drug latrunculin B (LatB); Left: WT and *sine1-1*; Right: WT and *sine2-1*. Symbols denote statistical significance as determined by Student’s t-test, with P<0.001. *: *sine1-1*, ABA+LatB vs. *sine1-1*, ABA only; ǂ: *sine2-1*, ABA+LatB vs. *sine2-1*, ABA only. (B) Buffers with and without 20 µM ABA and 10 µM of the F-actin stabilizing drug jasplakinolide (JK); Left: WT and *sine1-1*; Right: WT and *sine2-1*. P<0.001. *: specified lines vs. *sine1-1*, ABA only; ǂ: *sine1-1*, JK only vs. WT, JK only; ¥: *sine1-1*, ABA+JK vs. WT, ABA+JK. All data are mean values ± SE from three independent experiments.

Both under OB and OB + JK, stomata remained open in WT (Fig. 6B). JK inhibited ABA-induced closure in WT, as previously reported, indicating that actin depolymerization is necessary for ABA-induced stomatal closure (MacRobbie and Kurup, 2007; Li et al., 2014). OB alone resulted in sustained stomatal opening in *sine1-1* mutants. JK treatment in the absence of ABA actually led to stomatal closure in *sine1-1* mutants, as did the combination of JK and ABA (Fig. 6B, left panel). In contrast, JK did not induce stomatal closure in *sine2-1* and did not rescue the *sine2-1* defect in ABA-induced stomatal closure (Fig. 5A, right panel). To account for any differences seen in starting aperture size, the percent of stomatal closure was calculated (Supplemental Table S3) and the results are similar to those described above.

Together, these data indicate that actin depolymerization rescues the defect in ABA-induced stomatal closure that is caused by the loss of SINE1 or SINE2 and that, in the absence of SINE1, JK-induced actin stabilization and polymerization can mimic the effect of ABA.

## Discussion

We have shown here that the two related plant KASH proteins SINE1 and SINE2 play similar, yet distinguishable, roles in stomatal dynamics in response to light, dark, and ABA. We have previously reported that, in leaves, SINE1 is expressed specifically in guard cells and in the guard cell developmental lineage, while SINE2 is expressed predominantly in leaf epidermal and mesophyll cells, and only weakly detected in mature guard cells (Zhou et al., 2014). GFP-fusion proteins of SINE1 and SINE2 decorate the nuclear envelope, as expected based on their described KASH-protein function, but SINE1 is also detected in guard cells and mature root cells in a filamentous pattern that resembles actin and can be disassembled by actin-depolymerizing drugs (Zhou et al., 2014). In *sine1* mutant lines it was observed that the guard cell nuclei, which are typically arranged opposite each other in the center of the paired guard cells, are shifted from this position. This cellular phenotype was recapitulated in LatB treated wildtype guard cells, suggesting actin involvement, but was not found in *sine2* mutants. Based on these cell-biological data, we had hypothesized a function for SINE1, but not necessarily for SINE2, in guard cell biology.

Interestingly, in most bioassays applied here, *sine1* and *sine2* mutants showed similar guard-cell related phenotypes, which were also recapitulated by the double mutant, thus suggesting that the two proteins act in a shared pathway required for wildtype-like guard cell function. Loss of either SINE1 or SINE2 greatly diminishes stomatal opening in response to light, as well as stomatal closing in response to dark or ABA and significantly reduces the dynamic range of stomatal apertures between night and midday (Figs. 1 and 2). The lack of responsiveness to light for stomatal opening could be compensated in both single and double mutants through addition of external potassium and a low concentration of calcium. Both ions have been shown to play roles during light-induced stomatal opening (Hao et al., 2012; Wang et al., 2014), thus suggesting that the mutants might be hyposensitive to the external application of these ions (Fig. 1B).

During stomatal closure, ABA acts through Ca^2+^-dependent and Ca^2+^-independent signaling events (Wang et al., 2013). ABA increases both the Ca^2+^ entry at the plasma membrane and the internal Ca^2+^ release resulting in Ca^2+^ oscillations (Gilroy et al., 1991; Allen et al., 1999; Grabov and Blatt, 1999; Hamilton et al., 2000; Schroeder et al., 2001; Jiang et al., 2014; An et al., 2016). The increase in cellular Ca^2+^ is connected to ROS, inasmuch as a mutant of the NADPH oxidase which impairs ROS production also affects Ca^2+^ channel activation by ABA (Kwak et al., 2003). Conversely, intracellular Ca^2+^ activates the NADPH oxidase (Ogasawara et al., 2008), suggesting a positive feedback loop. This results ultimately in the activation of K^+^ outward channels and slow and fast ion channels (SLAC/ALMT), as well as the inactivation of K^+^ inward channels such as KAT1 (Jezek and Blatt, 2017). We therefore tested if application of the ROS H_2_O_2_ and/or Ca^2+^ could rescue the *sine* mutant stomatal closure phenotype. While H_2_O_2_ alone had no or a minimal effect, Ca^2+^ partially rescued the impaired stomatal closure and the combination of Ca^2+^ and ROS rescued the mutants to wildtype level (Fig. 4). Consistent with this finding, internal ROS increase after ABA exposure is still occurring in *sine1-1* and *sine2-1* (Supplemental Fig 7A and B). In contrast, Fura-2 staining suggests that early internal Ca^2+^ fluctuations after exposure to ABA are dampened in *sine1-1* and *sine2-1* (Supplemental Fig. 7C and D). Together, this suggests that, within ABA signaling, SINE1 and SINE2 act upstream of Ca^2+^, and that ROS exposure might intensify the Ca^2+^-based rescue, possibly through the described effect of ROS on activating the Ca^2+^ channels (Kwak et al., 2003; Wang et al., 2013; Jezek and Blatt, 2017).

Mutants with defects in stomatal regulation often show drought susceptibility phenotypes, due to their inability to fully close stomata and thus lose an excess of water through evaporation (Kang et al., 2002; Zhao et al., 2016). When we tested potted *sine* single and double mutants for increased drought susceptibility after withholding watering, or drought recovery defects after re-watering, we found no significant difference from wildtype plants (data not shown). Similarly, when detached rosette leaves were exposed to room air and monitored for fresh-weight loss, no difference from wildtype leaves was observed (Fig. 3A). In light of the results of the light/dark assay (Fig. 1A) as well as the impairment in both light-induced opening and ABA or dark-induced closing of stomata (Figs. 1B, 1C, and 2A), we argued that mutant plants and detached mutant leaves might be less subject to evaporation because, on average, stomata neither fully open nor fully close. This was substantiated by testing evaporation sensitivity after fully opening stomata by incubation in opening buffer. Now, indeed, mutant leaves wilted more rapidly, consistent with the observed defects in stomatal closure after this treatment (Fig. 3).

When the two lines that constitutively express SINE1 or SINE2 under control of the 35S promoter were added to this assay (Supplemental Fig. 3), we noted that both lines showed a fresh-weight loss indistinguishable from wildtype plants. This is not surprising in case of SINE1, given that this line showed no stomatal defects. Constitutive expression of SINE2, however, led to defects both in the light/dark assay and during ABA-induced stomatal closure (Fig. 5C and D) that would be consistent with a *sine*-mutant-like hypersensitivity to evaporation. To assess whether constitutive expression of SINE2 leads to additional - possibly compensatory - phenotypes, we calculated stomatal density and stomatal index of fully developed rosette leaves (Supplemental Fig. S6). Indeed, constitutive SINE2 expression leads to a reduction of both stomatal density and stomatal index. No such reduction was observed in any other line tested. The reduced number of stomata might compensate for the closing defects in this line, and thus result in wild-type-like evaporation within the level of resolution of our assay. Notably, this unique phenotype suggests that the SINE gene family not only acts in stomatal function but might also play a role during stomatal development. Only SINE1 is expressed throughout the guard cell developmental lineage, and mis-expression of SINE2 might thus highlight a yet unexplored role for SINE1 in the guard-cell developmental program.

It was noted that the *sine1-1 sine2-1* stomata respond differently from the *sine1-1* and *sine2-1* single mutants in the light/dark assay (Fig. 1A), but not in any of the subsequent assays. One notable difference between the light/dark assay and all other assays is that the former was performed on potted plants, using leaf imprints, while all other assays used detached leaves during treatments and epidermal peels for imaging. We thus argued that a whole plant phenotype might exist specifically in the double mutant that compensates for any stomatal dynamic defects seen in the light/dark assay. Indeed, we observed abnormalities in root architecture solely in the double mutant. As SINE1 and SINE2 are highly expressed in the seedling root (Zhou et al., 2014), we investigated primary root (PR) and lateral root (LR) characteristics (Supplemental Fig. S8). Of the four parameters tested, three showed significant differences in only the *sine1-1 sine2-1* double mutant: decreased PR length, number of LR, and LR density. No differences were observed in LR length. While it is currently unknown if this root morphology phenotype accounts for the behavior of the *sine1-1 sine2-1* mutant in the light/dark assay, these data demonstrate that the possibility of additional phenotypes unique to the double mutant have to be taken into account when interpreting these data.

Significantly less is known about the signal transduction pathway that triggers stomatal closure after exposure to dark. It is generally assumed, however, that many steps are shared with the ABA pathway (Jezek and Blatt, 2017). For example, mutants in the PYR/PYL/RCAR ABA receptors are also deficient in their stomatal response to darkness (Merilo et al., 2013). Similarly, diurnal rhythmicity is closely linked to the increase of guard cell ABA during the night - based both on de novo synthesis and import from the apoplast - and depletion of ABA levels during the day (Daszkowska-Golec and Szarejko, 2013). Together, these known connections are all consistent with the *sine* mutant phenotypes observed here, and suggest a primary role for SINE1 and SINE2 in a step downstream of ABA, but upstream of calcium, and downstream of some aspect of ROS enhancement of calcium-mediated guard cell closure.

A plethora of known effects of actin dynamics on stomatal aperture regulation (Kim et al.,1995; Gao et al., 2008; Gao et al., 2009; Higaki et al. 2010; Eun et al., 2001; MacRobbie and Kurup, 2007; Lemichez et al., 2001; Jiang et al., 2012; Li et al., 2014; Zhao et al., 2011), together with SINE1’s association with F-actin in guard cells made us speculate whether the *sine* mutant phenotypes are related to disturbances in guard cell actin dynamics. It has been shown previously that the inhibitor of F-actin assembly, latrunculin B (LatB) and the F-actin stabilizer, jaspakinolide (JK) have opposite effects on ABA-based stomatal closure (MacRobbie and Kurup, 2007). While LatB mildy enhanced closure in the presence of ABA, JK inhibited it. Our data recapitulate in Arabidopsis these effects reported for *Commelina communis* (MacRobbie and Kurup, 2007). Both *sine1* and *sine2* mutant phenotypes showed a clear interaction with the actin drugs. While loss of either SINE1 or SINE2 inhibited ABA-induced closure, co-incubation with LatB rescued this inhibition to a large degree (Fig. 6A). A hypothesis consistent with this finding is that SINE1 and SINE2 are connected to actin turnover during the transition from radial to longitudinal arrays that accompanies stomatal closure. In the presence of LatB, this activity would not be required. JK inhibited guard cell closure in both wildtype and the *sine2-1* mutant, consistent with the assumption that actin de-polymerization is a required step. Surprisingly however, JK alone, in the absence of ABA, was able to trigger stomatal closure in the *sine1-1* mutant (Fig. 6B). One scenario to explain this finding is that SINE1 might be required for an additional step involved in stabilizing F-actin, downstream of ABA, and that this step is also required for closing. In wildtype guard cells, this would be accomplished through an ABA-triggered involvement of SINE1, and JK has therefore no further effect. However, this step would be inhibited in the absence of SINE1, and could thus be rescued by JK alone, mimicking the ABA response. Clearly, more studies will be required to verify these proposed actin-related functionalities of the two proteins, but the data already show (1) that there is indeed an interaction between SINE1/2 function and actin worth further investigating, and (2) that while SINE1 and SINE2 have many overlapping functions, there are also critical differences, as revealed by the JK data. Future studies will have to focus on the real-time analysis of actin dynamics in the different mutant backgrounds and under the different treatments and the investigation of genetic interactions between *sine* mutants and the reported mutants of actin-modulating proteins involved in guard cell regulation.

Together, we have shown that the two plant KASH proteins SINE1 and SINE2 function in stomatal aperture regulation in a variety of scenarios that involve light, dark, and ABA. We propose that they act downstream of ABA, and upstream of calcium and actin. While there is a large degree of functional overlap between the two proteins, there are also critical differences, and their further analysis might shed light on the role of their intriguingly different expression patterns. This study reveals an unanticipated connection between stomatal regulation and a class of nuclear envelope proteins known to be involved in nuclear anchoring and positioning, and adds two novel players to the ever more complex world of guard cell biology (Albert et al., 2017). Addressing the connection between the phenotypes described here and the cellular role of SINE1 in guard cell nuclear positioning will likely be one of the more groundbreaking avenues of further study.

## Materials and methods

### Plant material

*Arabidopsis thaliana* (ecotype Col-0) was grown at 25°C in soil under 8-h light and 16-h dark conditions. For all assays, leaves were collected from 6-8 week-old Arabidopsis plants grown under these conditions. The *fls2-1* mutant has been reported previously (Uddin et al., 2017). *sine1-1* (SALK_018239C), *sine2-1* (CS801355), and *sine2-2* (CS1006876) were obtained from the Arabidopsis Biological Resource Center while *sine1-3* (GK-485E08-019738) was obtained from GABI-Kat. All SINE1 and SINE2 lines used here were previously reported (Zhou et al., 2014).

### Stomatal aperture measurements

Stomatal bioassays were performed by detaching the youngest, fully expanded rosette leaves of 6-8 week-old plants grown under short-day conditions (8 hours light; 16 hours dark).

For the light/dark assay, Duro super glue (Duro, item #1400336) was applied to a glass slide and the abaxial side of a leaf was pressed into the glue to create an imprint. Imprints were taken two hours prior to the chamber lights turning on and every two hours until two hours after the lights were turned off. The imprints were allowed to dry and subsequently imaged to obtain stomatal aperture measurements.

All other stomatal assays involved placing leaves in a petri dish abaxial side up with opening buffer (OB) containing 10 mM MES, 20 µM CaCl_2_, 50 mM KCl, and 1% sucrose at pH 6.15 for 3 h under constant light, leaves remained whole until designated time points at which abaxial epidermal strips were carefully peeled and imaged using a confocal microscope (An et al., 2016; Eclipse C90i; Nikon). For some of the experiments, OB base containing 10 mM MES and 1% sucrose at pH 6.15 was used to test stomatal dynamics with and without addition of 20 µM CaCl_2_ and/or 50 mM KCl. Stomatal closing assays were performed immediately after the opening assays, in which leaves were transferred to closing buffer containing 10 mM MES at pH 6.15 with or without the following treatments, as indicated: 20 µM ABA, 2 mM CaCl_2_, 0.5 mM H_2_O_2_, or 5 µM flg22 (Kang et al., 2002; Zhang et al., 2008b; Zhao et al., 2011). Leaves were placed in darkness to induce stomatal closure for 3 h when mentioned. NIS-Elements software was used for stomatal aperture measurements.

### Transpiration assay

To monitor water loss, five to six fully expanded leaves were detached from each plant at similar developmental stages (sixth to ninth true rosette leaves). For the “air only” assay, leaves were placed adaxial side up in an open petri dish under constant light on a laboratory bench and leaf weights were recorded every 30 minutes (Kang et al., 2002). For all other assays in Fig. 2, leaves were placed adaxial side up in either OB, OB base, OB base with 20 µM CaCl_2_, or OB base with 50 mM KCl for 3 h to induce opening. Leaves were then dried briefly on “Kim wipes” and placed adaxial side up in dry petri dishes for an additional 3 h under constant light. Leaf weights were recorded every 30 minutes. For the transpiration assay in Supplemental Fig. 2, leaves were placed abaxial side up in OB and then kept abaxial side up throughout the duration of the assay on paper towels. Leaves were weighed individually.

### Confocal microscopy and florescence intensity measurements

For the imaging and quantification shown in Figs. 5A and 5B, 6-8 week-old Arabidopsis leaves were imaged using a confocal microscope (Eclipse C90i; Nikon). Image settings were established first for the highest expressing line (35S:GFP-SINE1 in WT), to obtain a clearly visible, but not overexposed, GFP fluorescence signal at the nuclear envelope. These settings were then applied for imaging all other samples: a medium pinhole with a gain setting of 7.35 and the 488-nm laser set at 15% power. All images were taken at room temperature with a Plan Flour 40x oil objective (numerical aperture of 1.3, Nikon). NIS-Elements software was used to quantify fluorescence by drawing a region of interest around individual guard cell nuclei.

### ROS and calcium production assays

Detection of ROS in stomata was performed as described previously (Li et al. 2014). Whole leaves were incubated in OB adaxial side up for 3h under constant light. Leaves with open stomata were incubated in MES buffer pH 6.15 containing 50 µM of H_2_DCF-DA in the dark for 15 min and then washed with water. The leaves were then transferred to CB containing 20 µM ABA for 15, 30, 60 or 120 min. At the indicated time points, abaxial epidermal strips were peeled from the leaves for ROS detection by confocal microscopy with a setting of 488 nm excitation and 525 nm emission. The experiments were repeated four times with at least 70 stomata for each time point.

Detection of Ca^2+^ in stomata was performed using the Fura-2 AM dye (Sigma Aldrich, CAS 108964-32-5) (Jiang et al., 2014). Epidermal peels were floated in 10mM MES-TRIS (pH 6.1) buffer containing 1 µM Fura-2 AM and kept at 4°C in the dark for two hours. The Fura-2 dye was then washed out and peels were placed back in opening buffer for one hour at RT. ABA was added and time-lapse imaging of stomata was performed using confocal microscopy at specified time intervals.

### Immunoblotting

*N. benthamiana* leaves were collected and ground in liquid nitrogen into powder, and protein extractions were performed at 4°C. 1 ml radioimmunoprecipitation (RIPA) buffer was used to extract 500 µl of plant tissue, as described previously (Zhou et al., 2014). After three washes in RIPA buffer, samples were separated using 10% SDS-PAGE, transferred to polyvinylidene difluride membranes (Bio-Rad Laboratories), and detected with a mouse anti-GFP (1:2000; 632569; Takara Bio Inc.) or a mouse anti-tubulin (1:2000; 078K4842; Sigma-Aldrich) antibody. Membranes were imaged using an Odyssey Clx Imaging system and fluorescence was quantified using Image Studio software (LI-COR, inc).

### Yeast Strains and Manipulations

All work with yeast was done using *Saccharomyces cerevisiae* strain NMY51:MATahis3D200 trp1-901 leu2-3,112 ade2 LYS2::(lexAop)4-HIS3 ura3::(lexAop)8-lacZ ade2::(lexAop)8-ADE2 GAL4 obtained from the DUAL membrane starter kit N (P01201-P01229). Yeast cells were grown using standard microbial techniques and media (Lentze and Auerbach, 2008). Media designations are as follows: YPAD is Yeast Extract plus Adenine medium, Peptone, and Glucose; SD is Synthetic Defined dropout (SD-drop-out) medium. Minimal dropout media are designated by the constituent that is omitted (e.g. -leu –trp –his –ade medium lacks leucine, tryptophan, histidine, and adenine). Recombinant plasmid DNA constructs were introduced into NMY51 by LiOAc-mediated transformation as described (Gietz and Schiestl, 2007).

### Statistics

The number of stomata analyzed for each line, in all figures, is ≥80, unless otherwise stated. Error bars represent the standard deviation of means. Asterisks or symbols denote statistical significance after Student’s t-test as indicated.

## Supplemental Material

The following materials are available in the online version of this article.

**Supplemental Figure 1.** Stomatal opening in *sine* mutants prior to exogenous application of ABA.

**Supplemental Figure 2.** Stomatal closure in response to ABA for *sine1-3* and *sine2-2* mutants.

**Supplemental Figure 3.** Transpiration rates of individual leaves after induced stomatal opening.

**Supplemental Figure 4.** Protein blot analysis of transgenic Arabidopsis plants expressing GFP-SINE1 and GFP-SINE2.

**Supplemental Figure 5.** Interactions between SINE1 and SINE2 proteins in the membrane yeast two-hybrid system.

**Supplemental Figure 6.** Stomatal density and stomatal index of fully developed rosette leaves.

**Supplemental Figure 7.** ROS production and calcium monitoring in *sine* mutants.

**Supplemental Figure 8.** Root morphology of *sine* mutants.

**Supplemental Table S1.** Comparison of stomatal closure assays.

**Supplemental Table S2.** Percent stomatal closure for ABA and light-dark assays.

**Supplemental Table S3.** Percent stomatal closure for cytoskeleton drug treatment assays.

## Acknowledgements

This work was funded by a National Science Foundation grant to I.M. (NSF-1613501). We are grateful to Dr. Sarah Assmann (Penn State University) for critical reading of the first version of this manuscript. We would like to thank Andrew Kirkpatrick and all members of the Meier lab for many fruitful discussions throughout this work. We thank Dr. Xiao Zhou (Caribou Biosciences) for constructing the 35S-driven SINE1 and SINE2 expressing Arabidopsis lines while working in the Meier lab and Dr. David Mackey (OSU) for the *fls2-1* mutant.

## Notes

Funding: This work was supported by a grant from the National Science Foundation to I.M. (NSF-1613501).

