## Supplemental Figures and Tables for "Establishing the role of SINE proteins in regulating stomatal dynamics in Arabidopsis thaliana"

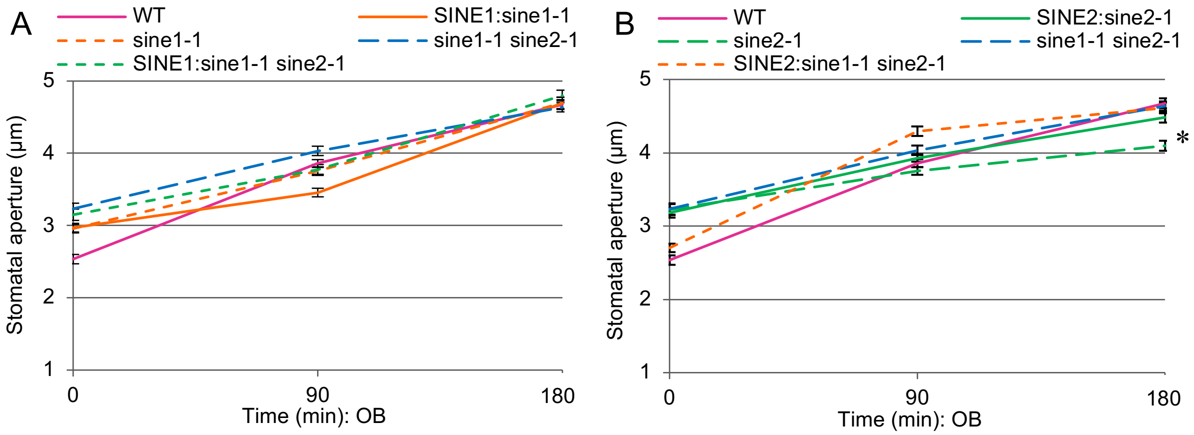


**Supplemental Figure 1: Stomatal opening in *sine* mutants prior to exogenous application of ABA.** Whole leaves were placed in opening buffer for 3 h under constant light at the end of a night cycle, epidermal peels were mounted every 90 min, and stomatal apertures were measured. (A) Lines including the *sine1-1* T-DNA insertion. (B) Lines including the *sine2-1* T-DNA insertion. Data from (A) and (B) obtained from one experiment and split into two panels for clarity. Data are mean values ± SE from three independent experiments. * denotes statistically significant difference as determined by Student's t-test, with $P<0.001$, between WT and *sine2-1*.

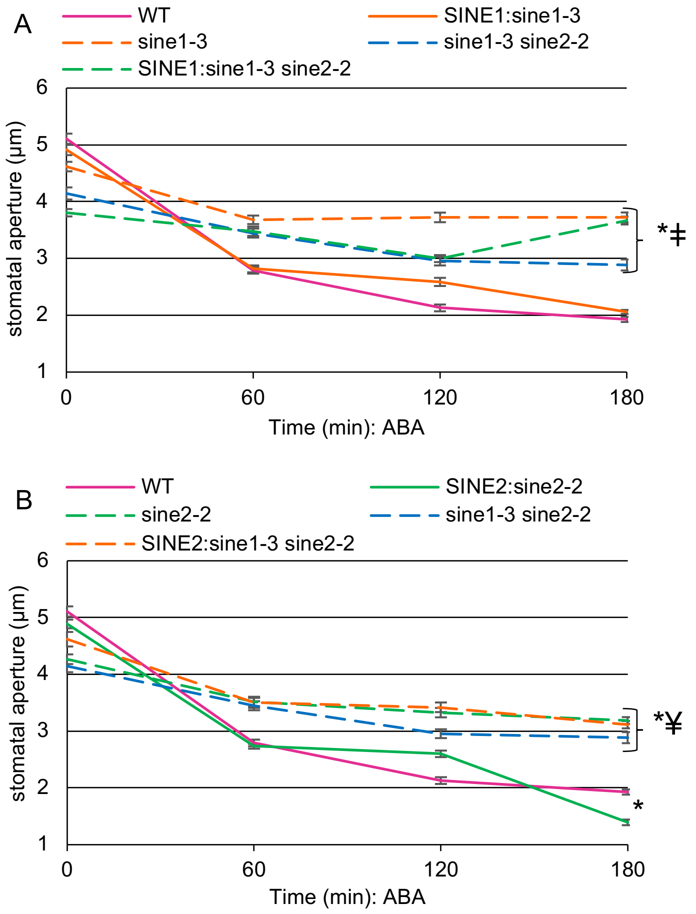


**Supplemental Figure 2: Stomatal closure in response to ABA for *sine1-3* and *sine2-2* mutants.** Stomatal opening and closing assays were used here as described in methods utilizing ABA to induce closure. (A) stomatal aperture measurement of SINE1:*sine1-3*, *sine1-3*, *sine1-3 sine2-2*, and SINE1:*sine1-3 sine2-2* (B) SINE2:*sine2-2*, *sine2-2*, *sine1-3 sine2-2*, and SINE2:*sine1-3 sine2-2*. Data obtained from one experiment and split into two panels for clarity. All data are mean values ± SE from three independent experiments. Symbols denote statistically significant differences, with P<0.001. *: specified lines vs. WT; ‡: specified lines vs. SINE1:*sine1-1*; ¥: specified lines vs. SINE2:*sine2-1*.

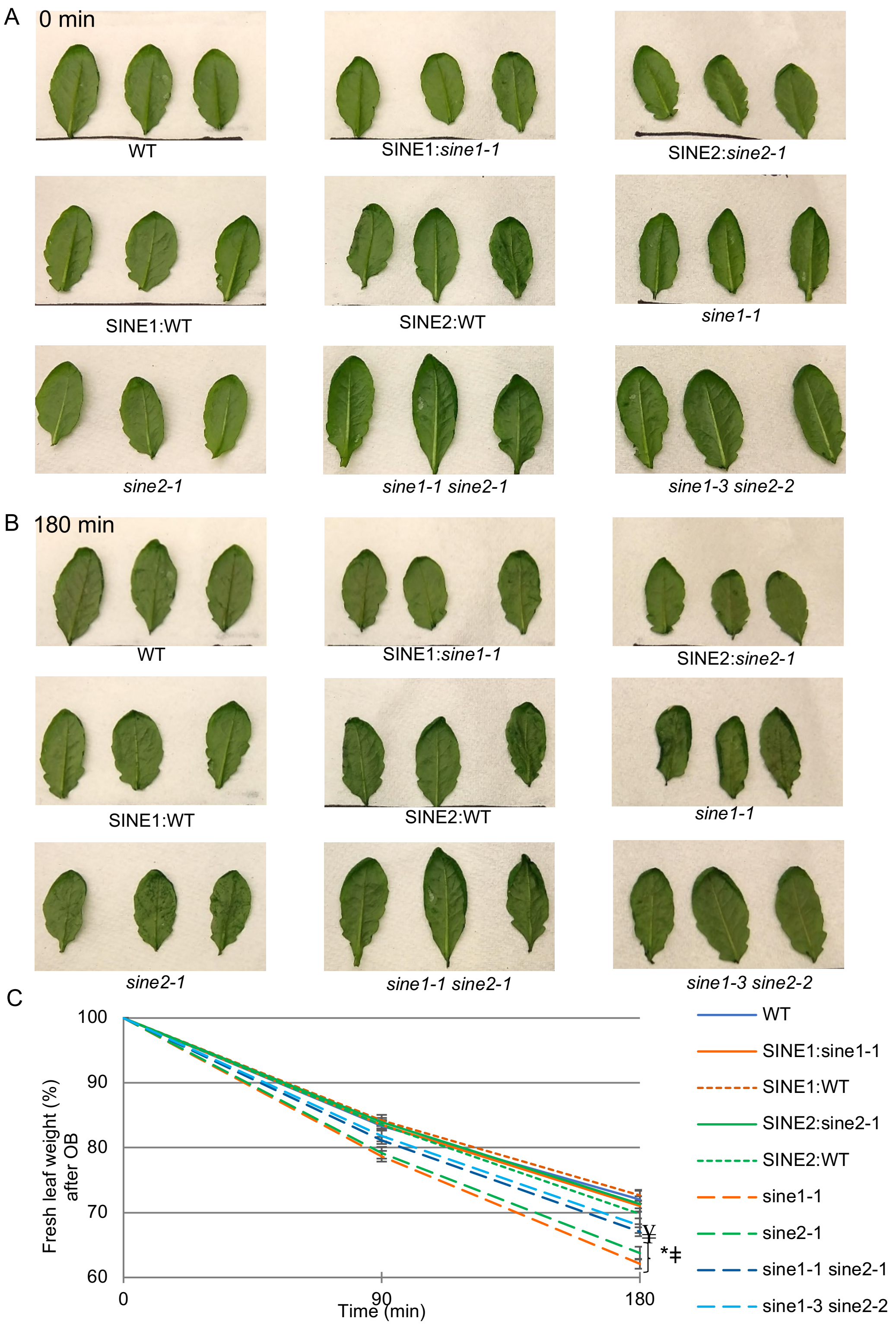


**Supplemental Figure 3: Transpiration rates of individual leaves after induced stomatal opening.** Rosette leaves were taken from 6-8 week old short day plants at similar developmental stages for each of the lines depicted and kept adaxial side up. Fresh leaves were placed in opening buffer 3 h under constant light, blotted dry, and transferred to a paper towel. Individual leaves were weighed immediately after drying and every 90 min. thereafter. (A) Representative images from one experiment at 0 min. (B) Representative images from the same experiment as in (A), taken at 180 min. (C) Quantification of leaf weight. Data are mean values ± SE from three independent experiments. Symbols denote statistically significant differences. *: P<0.001, specified lines vs. WT, SINE1:*sine1-1*, SINE1:WT, and SINE2:*sine2-1*; ǂ: P≤0.01 specified lines vs. SINE2:WT; ¥: P≤0.01 *sine1-1 sine2-1* vs. WT, SINE1:*sine1-1*, SINE1:WT, and SINE2:*sine2-1*.

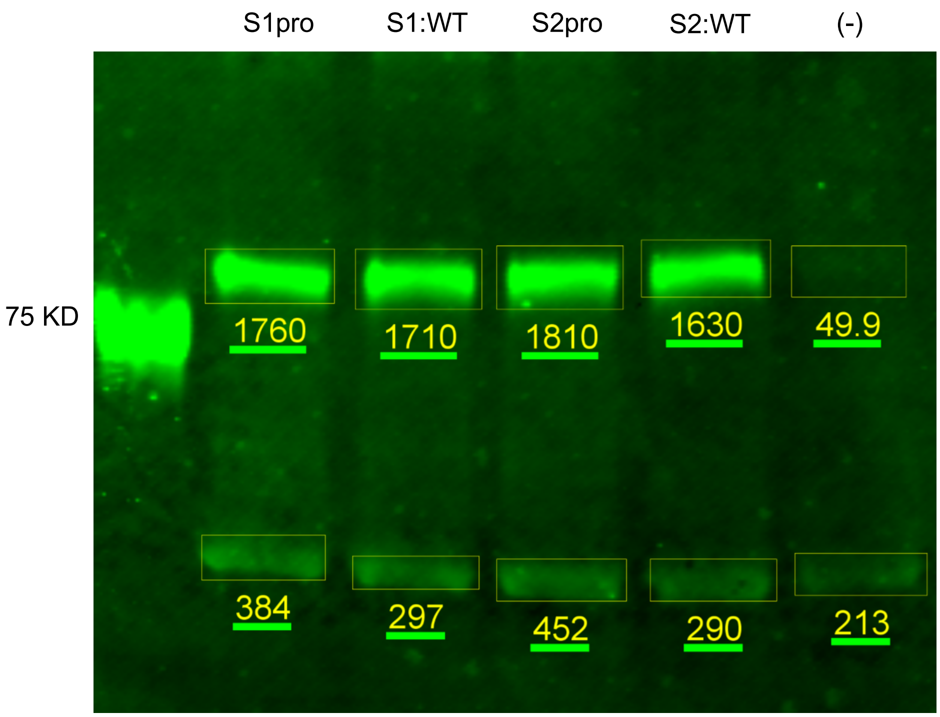


**Supplemental Figure 4: Protein blot analysis of transgenic Arabidopsis plants expressing GFP-SINE1 and GFP-SINE2.** Total protein was isolated from either rosette leaves or whole seedlings four independent times and separated using SDS-PAGE. Representative protein blot shown utilized whole seedlings. Top row representing, from left to right, $SINE1_{pro}$:GFP-SINE1 in *sine1-1* (S1pro), 35S:GFP-SINE1 in WT (S1:WT), $SINE2_{pro}$:GFP-SINE2 in *sine2-1* (S2pro), 35S:GFP-SINE2 in WT (S2:WT), and WT (-). Bottom row represents α-tubulin as a loading control. Protein blots were probed using Clontech mouse anti-GFP and IRDye® 800CW goat anti-mouse IgG and then imaged and quantified using Odyssey CLx from LI-COR.

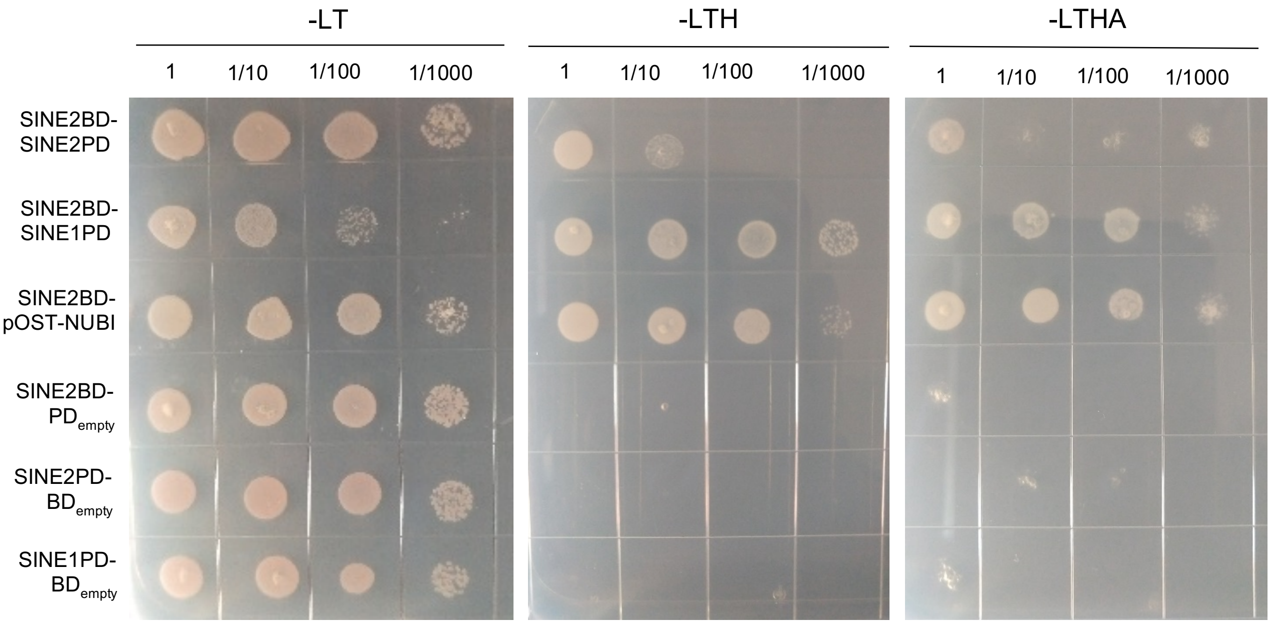


**Supplemental Figure 5: Interactions between SINE1 and SINE2 proteins in the membrane yeast two-hybrid system.** SINE1 and SINE2 were introduced into empty bait (BD) and prey (PD) vectors to test for possible protein-protein interactions. SINE1-BD was found to be self-activating and was thus not used to test for protein interactions. Negative controls utilized double transformations of either SINE1 or SINE2 with empty vectors. Control plates contained synthetic dropout (SD) media lacking –leu –trp while test plates contained SD media lacking –leu –trp –his or –leu –trp –his –ade. Double yeast cultures were diluted as shown to test for interaction strength.

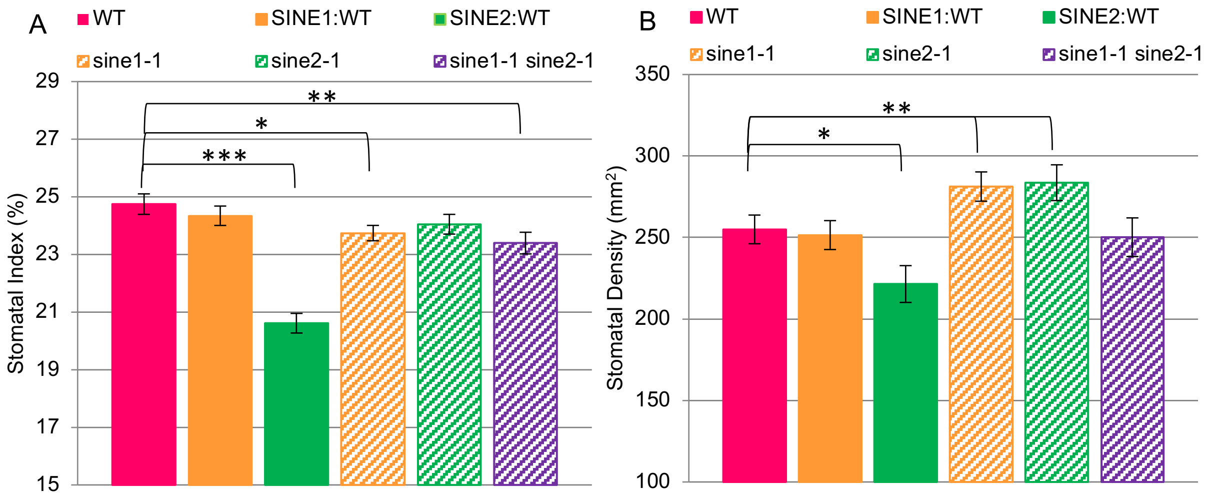


**Supplemental Figure 6: Stomatal density and stomatal index of fully developed rosette leaves.** Epidermal peels were used for quantification. Stomatal index (SI) determines the ratio of stomata to pavement cells as stomata are known to follow specific spacing rules between these two cell types: SI(%)= #stomata/(#stomata + #pavement cells)*100 (Casson et al., 2009). Stomatal density (SD) measures how many stomata are in a given area: SD= #stomata/mm2 (Casson et al., 2009). (A) Stomatal index. Asterisks denote statistically significant differences. *** P<0.001; ** P=0.01; * P=0.03;. (B) Stomatal density. * P=0.02; ** P<0.05. All data represent mean values ± SE of 8 individual plants and ≥ 600 stomata per transgenic line.

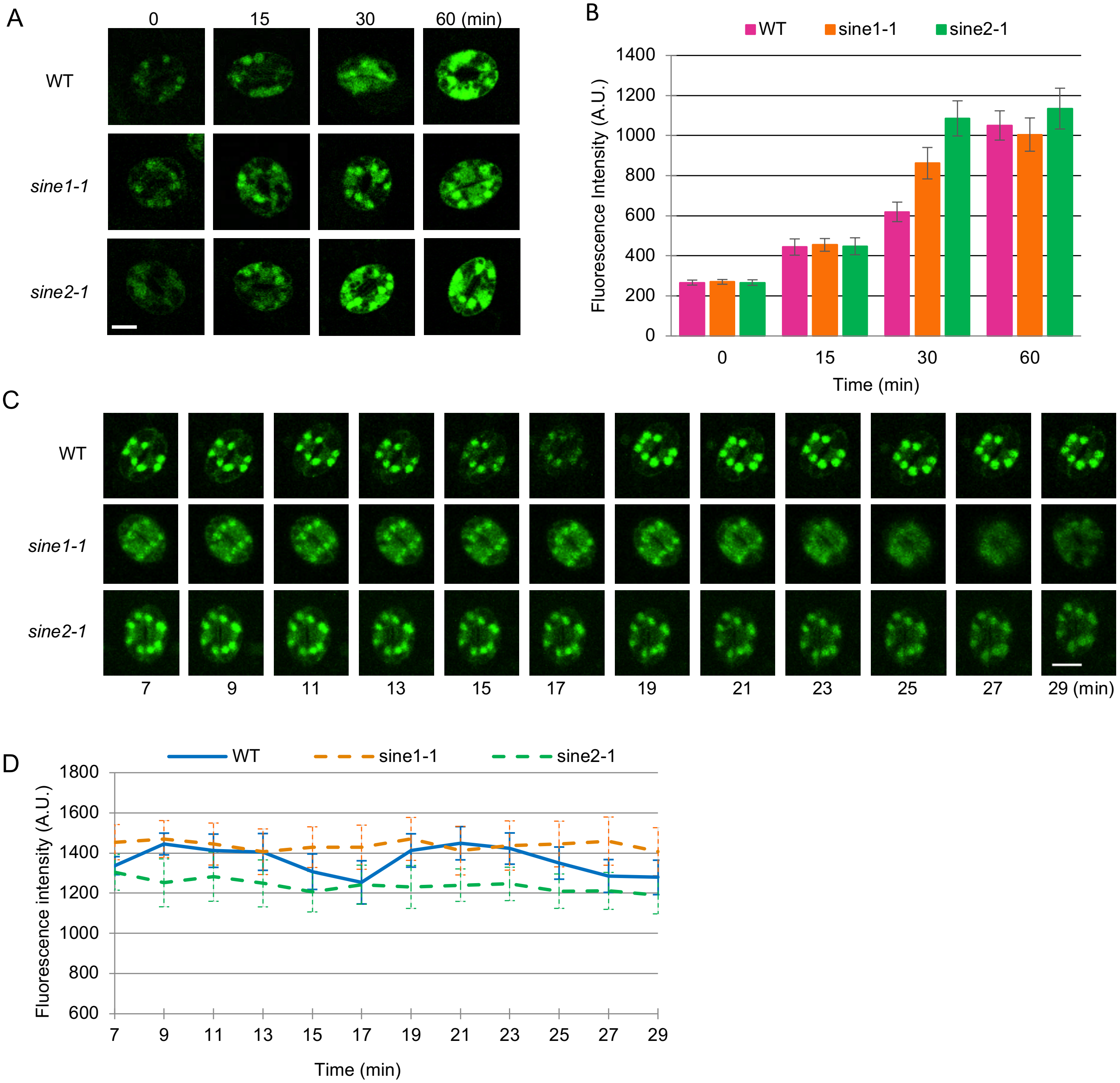


**Supplemental Figure 7: ROS production and calcium monitoring in *sine* mutants**. Stomatal opening and closing assays were used here as described in methods. (A) Confocal microscopy was used to take images of epidermal peels incubated in $H_2$DCF-DA dye, representative fluorescence images shown. The same laser and gain settings were used for all the images. Scale bar represents 10 µm. (B) Fluorescence intensities representing ROS levels in stomata of WT, *sine1-1,* and *sine2-1* during ABA treatment with four independent repeats and ≥ 70 stomata per line. (C) Confocal microscopy was used to take images of epidermal peels incubated in Fura-2 dye, representative fluorescence images shown. The same laser and gain settings were used for all the images. Scale bar represents 10 µm. (D) Fluorescence intensities representing calcium levels in stomata of WT, *sine1-1*, and *sine2-1* during ABA treatment with four independent repeats and ≥ 15 stomata per line. The data are presented as the mean values ± SE.

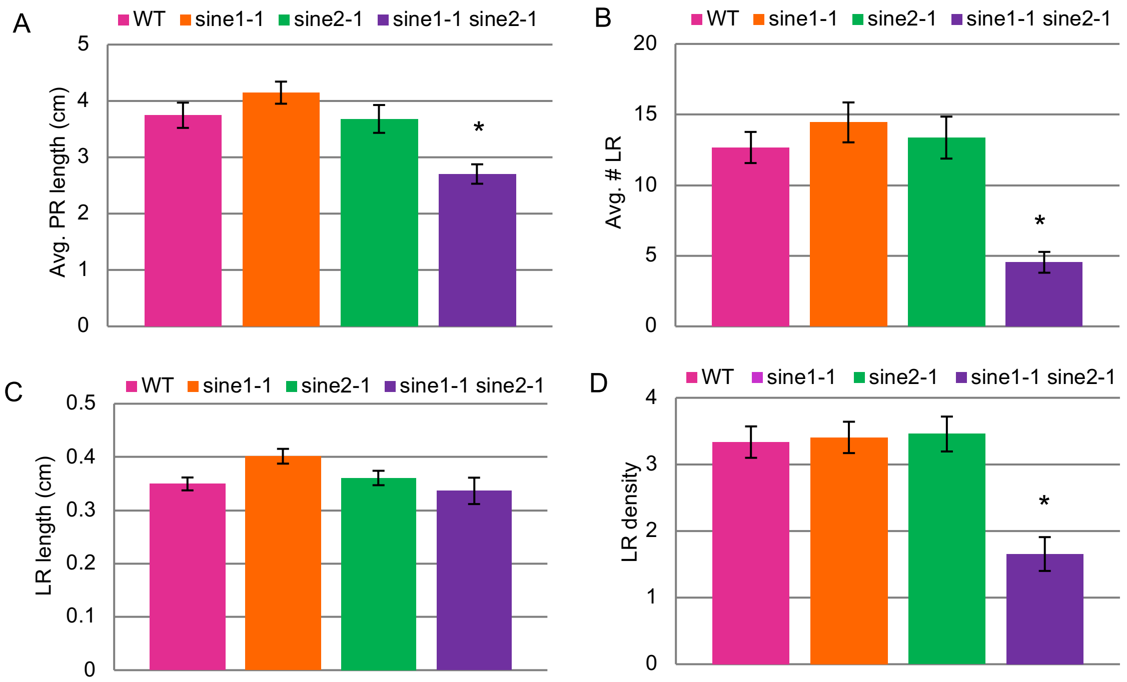


**Supplemental Figure 8: Root morphology of *sine* mutants.** Arabidopsis seeds were grown on MS media lacking sucrose for 11 days and root morphology was assessed using Image J. (A) Average primary root (PR) length. (B) Average number of lateral roots (LR). (C) LR density, as measured by (LR #/PR length). (D) average LR length. ≥27 seedlings were analyzed per line. Asterisk denotes statistically significant differences, with $P<0.001$ for *sine1-1 sine2-1* vs. WT.

**Supplemental Table 1: Comparison of stomatal closure assays.** Stomatal apertures at the end of ABA- (Fig. 2A), $H_2O_2$- (Fig. 4A-B), and darkness (Fig. 1C)-induced stomatal closure assays were directly compared to stomatal apertures at the end of the $CaCl_2$ (Fig. 4E)-induced stomatal closure assay.

| **180 min closure** | **WT** | | ***sine1-1*** | | ***sine2-1*** | | ***sine1-1 sine2-1*** | |
|---|---|---|---|---|---|---|---|---|
| | **Stomatal apertures ±SE** | **P-value** | **Stomatal apertures ±SE** | **P-value** | **Stomatal apertures ±SE** | **P-value** | **Stomatal apertures ±SE** | **P-value** |
| **$CaCl_2$ vs. ABA** | 2.11±0.05; 2.01±0.04 | =0.16 | 2.47±0.08; 3.79±0.07 | <0.001 | 2.53±0.08; 3.71±0.07 | <0.001 | 2.84±0.08; 3.48±0.05 | <0.001 |
| **$CaCl_2$ vs. $H_2O_2$** | 2.11±0.05; 2.18±0.05 | =0.33 | 2.47±0.08; 3.35±0.06 | <0.001 | 2.53±0.08; 3.09±0.06 | <0.001 | 2.84±0.08; 3.15±0.06 | =0.005 |
| **$CaCl_2$ vs. Darkness** | 2.11±0.05; 2.91±0.05 | <0.001 | 2.47±0.08; 4.13±0.06 | <0.001 | 2.53±0.08; 4.01±0.06 | <0.001 | 2.84±0.08; 4.02±0.06 | <0.001 |

**Supplemental Table 2: Percent stomatal closure for ABA and light-dark assays.** Stomatal closure assays reported here, converted from stomatal aperture values to percentages. Starting apertures are set to 100% and compared to apertures at the end of each assay, with smaller percentages meaning increased closure. Percent closure for the following assays: ABA-induced closure (Fig. 3A), $H_2O_2$-induced closure (Fig. 4A-B), $Ca^{2+}$-induced closure (Fig. 4C-D), $H_2O_2$ + $Ca^{2+}$-induced closure (Fig. 4E), and darkness-induced closure (Fig. 1C).

| Treatment | WT | SINE1: *sine1-1* | SINE2: *sine2-1* | *sine1-1* | *sine2-1* | *sine1-1 sine2-1* | SINE1: *sine1-1 sine2-1* | SINE2: *sine1-1 sine2-1* |
|---|---|---|---|---|---|---|---|---|
| **ABA** | 43% | 44% | 48% | 80% | 91% | 75% | 71% | 79% |
| **$H_2O_2$** | 45% | 49% | 54% | 83% | 83% | 76% | 79% | 75% |
| **$Ca^{2+}$** | 45% | 48% | 46% | 64% | 59% | 62% | 63% | 59% |
| **$H_2O_2$ + $Ca^{2+}$** | 67% | 60% | 62% | 70% | 67% | 67% | N/A | N/A |
| **Darkness** | 66% | N/A | N/A | 97% | 86% | 92% | N/A | N/A |

**Supplemental Table 3: Percent stomatal closure for cytoskeleton drug treatment assays.** Stomatal closure assays reported here, converted from stomatal aperture values to percentages. Starting apertures are set to 100% and compared to apertures at the end of each assay, with smaller percentages meaning increased closure. Percent closure for the following assays: LatB treatment (Fig. 6A) and JK treatment (Fig. 6B).

| Treatment | Fig. # | Grouped data | | Grouped data | |
|---|---|---|---|---|---|
| | | **WT** | ***sine1-1*** | **WT** | ***sine2-1*** |
| **OB** | 6A | 92% | 101% | 98% | 98% |
| **LatB** | | 101% | 98% | 91% | 93% |
| **ABA** | | 60% | 85% | 59% | 90% |
| **ABA + LatB** | | 54% | 66% | 60% | 65% |
| **OB** | 6B | 100% | 106% | 93% | 101% |
| **JK** | | 108% | 75% | 93% | 99% |
| **ABA** | | 53% | 103% | 54% | 96% |
| **ABA + JK** | | 93% | 84% | 87% | 93% |
